# The nuclear receptor NR4A1 restrains neutrophil granulocyte mediated brain damage in cerebral ischemia

**DOI:** 10.1101/2022.02.27.482146

**Authors:** Jan-Kolja Strecker, Marie Liebmann, Julian Revenstorff, Carolin Beuker, Antje Schmidt-Pogoda, Stephanie Hucke, Thomas Vogl, Johannes Roth, Christian Thomas, Tanja Kuhlmann, Heinz Wiendl, Luisa Klotz, Jens Minnerup

## Abstract

Immigration and activation of immune cells play a significant role in damage progression after ischemic stroke. It has been shown that the nuclear receptor NR4A1 exerts a crucial role within the inflammatory response of various immune diseases via regulating immune cell activation. In this study, we investigated the role of NR4A1 on the activation and recruitment of brain resident and peripheral immune cells after cerebral ischemia. Here, we show that NR4A1 mediates an anti-inflammatory and damage limiting effect after ischemic stroke through immigrating neutrophil granulocytes. Importantly, NR4A1-activation with its ligand Cytosporone-B improves functional outcome and diminishes brain damage. Therefore, modulation of NR4A1 is a promising therapeutic target in the treatment of stroke.

## Introduction

Even after decades of research efforts, stroke still is one of the main causes of death and disability worldwide (Benjamin et al., 2017). To date, only two approved specific therapeutic options are available to a minority of patients during a narrow time-window after stroke onset, intravenous thrombolysis and thrombectomy (Powers et al., 2018) (David and Donnan, 2009). Due to several contraindication only 20% of ischemic stroke patients receive these specific therapies. During the first days after the onset of ischemia the local inflammatory response significantly contributes to brain damage (Jayaraj et al., 2019). The stroke induced neuroinflammation particularly includes activation of resident micro- and astroglia as well as infiltration of circulating myeloid cells (Gelderblom et al., 2009; Beuker et al., 2021). However, substantial questions regarding the impact of different immune cells on brain injury remain unanswered. Hematogenous leukocytes, for example, show phase-dependent effects in stroke pathology and have been shown to exacerbate brain damage, mediate secondary neuronal damage but also contribute to tissue damage resolution during the chronic phase (Hu et al., 2012; Chernykh et al., 2016; Semerano et al., 2019). On the other hand, brain resident microglia has been shown to exert protective effects during acute ischemic injury (Szalay et al. 2016). Overall, the increasingly deeper understanding of the pathophysiology within these damage cascades opens up new therapeutic options.

Nuclear receptors represent a promising option for the therapeutic modulation of activation and infiltration of immune cells. Nuclear receptors are DNA-binding transcription factors regulating a plethora of biological programs in adaptation to cellular stress and in pathophysiology including cell proliferation, apoptosis, metabolism, ontogenesis and immune-cell activation (Mazaira et al., 2018, Wu et. Chen, 2018). The nuclear receptor NR4A1 (also known as NUR77, NAK-1, NGFI-B, GFRP1, TR3, N10) is an immediate early gene receptor and belongs to the group of nerve growth factor IB-like orphan nuclear receptors which are induced by diverse stressors. NR4A receptors have been implicated in the pathogenesis of various immune-mediated as well as non-immune-mediated diseases including psoriasis, diabetes, rheumatoid arthritis, atherosclerosis and malignancies (Volokakis et al. 2008; Huang et al., 2014; Xiong et al., 2020). NR4A1 acts as a key suppressor of pro-inflammatory cytokines such as IL-1β, IL-6, and CXCL-8 in monocytes, macrophages and T cells, among others via inhibition of NF-κB, (Hanna et al., 2011, 2012). Based on the regulation of NR4A1 in inflammatory processes as well as in different forms of cerebral injury (Liebmann et al., 2018; Wang et al., 2018), we hypothesized that NR4A1 might have a crucial role within the pathophysiology of cerebral ischemia.

With the use of different loss-of-function approaches in NR4A1-deficient, chimeric and cell-specific knock-out mice we were able to narrow the NR4A1-protective effects down to immigrating neutrophil granulocytes. We could show that NR4A1 limits activation as well as immigration of neutrophil granulocytes into the lesion after experimental stroke. Further, ligand mediated NR4A1-activation with Cytosporone-B (Zhang et al., 2008) resulted in improved functional outcome and reduced infarct size. In summary, these data suggest that the nuclear receptor NR4A1 has a key damage-limiting role within the pathophysiology of stroke particularly mediated by the activation of infiltrating neutrophil granulocytes. Targeting this rector could resemble a promising therapeutic target in stroke therapy.

## Methods

### Animals

All animal procedures were approved by the local governmental authorities (Landesamt für Natur, Umwelt und Verbraucherschutz, NRW, Germany, and Landesamt für Verbraucherschutz, Thüringen, Germany) and performed in accordance with local animal welfare regulations. All C57BL/6J, NR4A1-deficient (Lee et al., 1995) and Rag-1-deficient (Mombaerts et al., 1992) mice were maintained under pathogen free conditions on a 12:12 hour light-dark cycle period with access to food and water ad libitum. Adult male mice (10-16 weeks with C57Bl/6 background) were used in all experiments. Genotyping was performed according to Jax Mice genotyping protocols, via PCR using DNA extracted from ear punches.

### Bone marrow chimeras

Bone marrow chimeric mice were generated via transplantation of 0.5 x 10^7^ congenic bone marrow cells of either wildtype (C57Bl/6J, CD45.2) or NR4A1-deficient mice (CD45.2) into the vena coccygealis of sublethally irradiated (7.5 Gy) congenic wildtype recipient mice (C57Bl/6J, CD45.1). Proper chimerism was validated following 8 weeks of reconstitution by flow cytometry analysis for allelic CD45 variants within blood samples collected from the tail (CD45.1 vs. CD45.2). Animals with >88% CD45.2^+^ and <12% CD45.1^+^ leukocytes were used for subsequent experiments.

### Cytosporone-B treatment

Cytosporone-B or the corresponding placebo (16.5% DMSO in PBS) was applied to wildtype mice (13 mg/kg body weight, i.p.; Dai., et al., 2014) 3h or 12h after induction of reperfusion. Further, NR4A1-deficient mice received either Cytosporone-B or the solvent 3h after reperfusion.

### Adoptive T cell transfer

For preparation of spleen lymphocytes, two wildtype and two NR4A1-deficient mice (age 12-16 weeks) were sacrificed, the spleens separated and squeezed through a cell strainer (70 μm mesh). Cells were then washed by centrifugation (300 rpm, 5 min), resuspended in MACS-buffer (PBS with 1% FCS and 0.4% EDTA) and once again squeezed through a cell strainer (40 μm mesh). Cells were then washed in MACS-buffer und T cells were MACS-sorted from suspension using a Pan T Cell Isolation Kit (Miltenyi Biotec, Bergisch Gladbach, Germany). Subsequently, either sorted wildtype or NR4A1-deficient T cells (5×10^6^) were transferred intravenously to Rag-1-deficient mice (which lack T and B cells). Experimental cerebral ischemia was induced by the occlusion of the middle cerebral artery 24h after T cell transfer. After 24h of MCAO, sufficient T cell transfer was confirmed by flow cytometry analysis of CD3^+^-T cells in the blood.

### Middle cerebral artery occlusion

Occlusion of the middle cerebral artery (MCAO) was induced in mice under inhalation anesthesia (1.5% isoflurane in 30% O_2_/70% N_2_O) and maintenance of 37°C body temperature during the whole procedure. After midline neck incision, the left common artery and carotid bifurcation were exposed and the external and proximal left common arteries were ligated subsequently. Retrograde perfusion of the left common carotid artery was transiently interrupted using a microvascular clip (FE691; Aesculap, Tuttlingen, Germany). The common carotid artery was then incised using a micro-dissecting scissors, and a silicon-coated 8–0 nylon monofilament (701956PK5Re or 702056PK5Re, Doccol Corporation, Sharon, MA) was advanced into the middle cerebral artery. Following 30 minutes of MCA occlusion, the filament was retracted to allow reperfusion of the MCA.

### Functional testing

Neurological deficit score rating was performed in wildtype and NR4A1-deficient animals over the time-course of 3 days following MCAO: normal motor function (0); reduced grip of contralateral forelimb grip while tail pulled (1); flexion of torso and contralateral forelimb when mouse was lifted by the tail (2); circling to the contralateral side when mouse was held by the tail on a flat surface, but normal posture at rest (3); no spontaneous motor activity (4) or dead (5). For assessment of skilled motor performance, the rotarod test was used. In the rotarod test, mice were placed on a rotating cylinder (4–50 rpm acceleration in 210 s; TSE-Systems, Germany), and the time the animals remained on this cylinder was measured. Mice were given 3 training sessions before baseline testing. Mean duration of animals running on the rod was recorded 1 day prior and daily after MCAO for 3 consecutive days. For comparison of sensorimotor deficits the foot fault test was used. In the foot fault test, the animals were placed individually on an elevated 10-mm square wire mesh (total grid area of 40 cm x 40 cm) and subsequently videotaped while walking freely for 2 minutes. The number of foot faults as well as the total number of steps was counted. The percentage of foot faults was calculated as number of foot faults of the affected limb / number of total steps of the affected limb*100.

### Tissue collection and processing for histology

Six hours, 24h or 72h after MCAO, mice were perfused through the left ventricle with 30ml phosphate buffered saline (PBS) for FACS and mRNA-analyses or for 5 minutes with PBS followed by 4% paraformaldehyde solution for 10 minutes for histology tissue preparation. Mice were held under deep xylazine/ketamine anesthesia during perfusion. Brains were removed and transferred to PBS (flow cytometry analyses) or frozen on dry ice (mRNA analyses). For histological staining, brains were fixed in 4% paraformaldehyde overnight, immersed in 20% sucrose for three days, embedded in TissueTek^®^ frozen and stored at −80°C.

### Infarct volume assessment

Infarct volumes were retrieved by collecting 10 μm coronal cryosections in intervals of 300 μm starting at the rostral border of the infarction. Subsequently, slices were stained with 0.5% toluidine blue and dried in graded ethanol (1 min each) at concentrations of 50%, 80%, 96% and 100%. Digitized images were then measured, using ImageJ software, by an investigator blinded to genotype or treatment. Infarct volume was calculated by multiplying infarct area size by 300 μm. Edema compensation was applied using the formula (area of contralateral hemisphere / area of ipsilateral hemisphere)*infarct area. Infarct volumes ≤ 7 mm^3^ were considered as insufficient occlusion and therefore excluded (1x Cytosporone-B treatment group, 3h; 1x placebo group, 3h).

### Immunohistochemistry

Mounted coronal cryosections were rinsed in PBS for 3 times (5 min) and thereafter incubated in Blocking Reagent (Roche Diagnostics) for 15 minutes. We used the following antibodies for the murine sections: anti-GFAP (1:500, Dako, raised in mouse), anti-NeuN (1:150, Millipore, raised in mouse), anti-F4/80 (1:500, Serotec, raised in rat), anti-Laminin (1:100, Abcam, raised in rabbit), anti-Ly-6B.2 (1:100, clone 7/4, BioRad, raised in rat) and anti-Nur77 (NR4A1, 1:100, Abcam, raised in goat). NeuN and GFAP antibodies were visualized with anti-mouse-488 secondary antibody (1:150, 45 min, Life). Mouse IgG was visualized using an anti-mouse-594 antibody (1:150, 45 min, Life). F4/80 and Ly6B.2 antibodies were visualized with an anti-rat-488 antibody and NR4A1 was visualized with an anti-goat-594 antibody (1:100; 45 min, Jackson ImmunoResearch). Human brain sections were rehydrated and antigen demasking was done using EnVisionTM FLEX Target Retrieval Solution (Agilent Technologies, pH 9.0) under heat. After washing 3 times in PBS for 5 min, sections were blocked for 15 minutes in Blocking Reagent (Roche Diagnostics, room-temperature) and subsequently incubated with primary antibodies anti-Nur77 (NR4A1, 1:100, Abcam, raised in goat) and Neutrophil Elastase (1:100, Abcam, raised in rabbit, 4°C over night). Visualization of Neutrophil Elastase was done using an anti-rabbit-488 antibody (1:100, 45 min) and NR4A1 was either tagged with an anti-goat-594 antibody or made visible using 3,3’-Diaminobenzidin staining according to the manufactures protocol (Sigma-Aldrich). Fluorescence stainings were mounted with Vectashield Mounting Medium with DAPI (Vector). Images were taken with a Nikon Eclipse 80i fluorescence microscope (Nikon) and a Zeiss AxioVision Apotome (Carl Zeiss). For generation of heatmaps, traced areas of NR4A1-fluorescence signal profiles (tracked at ~0.5 mm bregma) were transferred to Adobe Illustrator and overlaid. Saturation of each animal’s occupied area was set to 1/n intensity adding up to 100% saturation in areas of full overlay within all n=11-12 animals.

### Real-Time Polymerase Chain Reaction

Primers used for expression studies were purchased from Qiagen (QuantiTect Assays, Hilden. Germany): *Ccl2* (QT00167832), *Cxcl1* (QT00115647), *Il6* (QT00098875), *Nfkbia* (QT00134421), *Nos2* (QT00100275), *Nr4a1* (QT00101017), *Pgk1* (QT00306558), *Ptgs2* (QT00165347), *Tnf* (QT00250999). Complementary DNA (cDNA) concentrations were measured by semi-quantitative real-time polymerase chain reactions (RT-PCR) using the BioRad CFX384 RT-PCR-System (Hercules, CA; USA) and SYBR-green fluorescence. Standard LightCycler conditions were used for the expression arrays, and all measurements were done in duplicates. Stored cDNA-libraries from previous investigations (12h, 18h, 24h, 36h and 168h) have been used for examination of *Nr4a1*-expression after MCAO.

### Expression Analysis

Gene expression was related to the individual expression of phosphoglycerate-kinase (*Pgk1*) as endogenous control. Expression analysis was performed using expression software tool REST-MCS V2 (Pfaffl et al., 2002). In cases when cDNA samples did not exceed the set RT-PCR threshold after 40 cycles (not determined), the respective ΔCT was equated with 40 for the further analysis.

### Human samples

Brain autopsy material of 2 patients suffering from acute ischemic stroke who died after stroke onset at the University Hospital of Münster, Germany was analyzed in this study. Research was conducted in accordance with the declaration of Helsinki and was approved by the local ethics committee (approval ID 2017-210-f-S). Samples were derived from brain regions mainly comprising the frontal cortex, striatum, and internal capsule, therefore contained in the vascular territory of the middle cerebral artery.

### Metabolic assay of HoxB8-immortalized granulocytes

Metabolic activity of HoxB8-immortalized wildtype or NR4A1-deficient granulocytes (Gran et al. 2018) were determined with XFe96 Extracellular Flux Analyzer (Agilent Technologies), as described previously (Liebmann et al., 2018). Following five days of differentiation, 200.000 HoxB8-immortalized granulocytes were cultured in XF Base Medium Minimal Dulbecco’s Modified Eagle’s Medium (Agilent Technologies) containing 10 mM glucose, 2 mM L-glutamine and 1 mM sodium pyruvate (all Merck). The oxygen consumption rate (OCR) and extracellular acidification rate (ECAR) were evaluated under basal conditions and in response to 2 μM oligomycin, 1 μM FCCP, 100 nM rotenone, plus 1 μM antimycin A (all Sigma-Aldrich). In advance, cells were treated either with or without 600 nM phorbol-12-myristat-13-acetat (PMA, Sigma-Aldrich) for 3 h and OCR and ECAR were determined in parallel. OCR and ECAR were analyzed using the Wave Desktop software (Agilent).

### Blinded assessment

Analysis of functional testing, infarct volume assessment and quantification of histological findings was performed in a blinded manner by two experienced medical technical assistants and reviewed by JKS.

### Antibodies and flow cytometry

Surface marker staining was performed as described (Klotz L, et al., 2009). Intranuclear staining was performed employing Foxp3/Transcription Factor Staining Buffer Set (eBioscience). All murine antibodies used are summarized in Table S1. Flow cytometric analysis was done on a Gallios Flow Cytometer or a CytoFLEX (both from Beckman Coulter) and results were analyzed with FlowJo (Tree Star) or Kaluza (Beckman Coulter) software.

### Statistical analysis

Statistical analysis was performed using GraphPad Prism version 8 (GraphPad Software, La Jolla, CA). Data were checked for normal distribution applying the Shapiro-Wilk normality test followed by group comparison using the Student’s *t* test or the Mann-Whitney test. Repeated measures were analyzed with 2way ANOVA and subsequent Bonferroni post-hoc test. Data are presented as mean ± SEM unless otherwise stated. A p value of <0.05 was considered significant.

## Results

### Stroke induces dynamic expression of NR4A1 within the ischemic brain hemisphere

Stroke induces a substantial inflammatory response within the CNS parenchyma accompanied by activation of brain resident immune cells and infiltration of hematogenous immune cells (Beuker et al., 2021). As the nuclear receptor NR4A1 has been shown to play a central role in various pathophysiological immune-mediated processes (Koenis et al., 2018; Pulakazhi et al., 2021), we have hypothesized that NR4A1 may also play a role during the progression of immune-mediated post-ischemic brain damage. As a first step, we evaluated *Nr4a1-mRNA* and NR4A1-protein expression after middle cerebral artery occlusion (MCAO) in C57BL/6 mice (30 min MCAO) over a time-course of 7 days post ischemia. Analysis of whole-brain lysates by qRT-PCR revealed a dynamic and time-dependent *Nr4a1*-expression during the acute (12h post MCAO) and the sub-acute phase (7 days post MCAO, Fig. 1A). Immunohistological staining confirmed that stroke results in a time- and lesion-dependent NR4A1-induction within the affected hemisphere 24h, 72h and 168h after MCAO (Fig. 1B). Co-localization studies revealed particularly neutrophil granulocytes, damaged neurons and endothel-associated cells as main NR4A1-expressing cell populations (Fig. 1C). Analysis of stroke patient brain slices confirmed neutrophil granulocytes and neurons as source for NR4A1 within the ischemic human brain (Fig. 1D). These data suggest that NR4A1 is mainly expressed by neutrophil granulocytes and neurons during the acute and the sub-acute phase of stroke.

**Figure 1.**
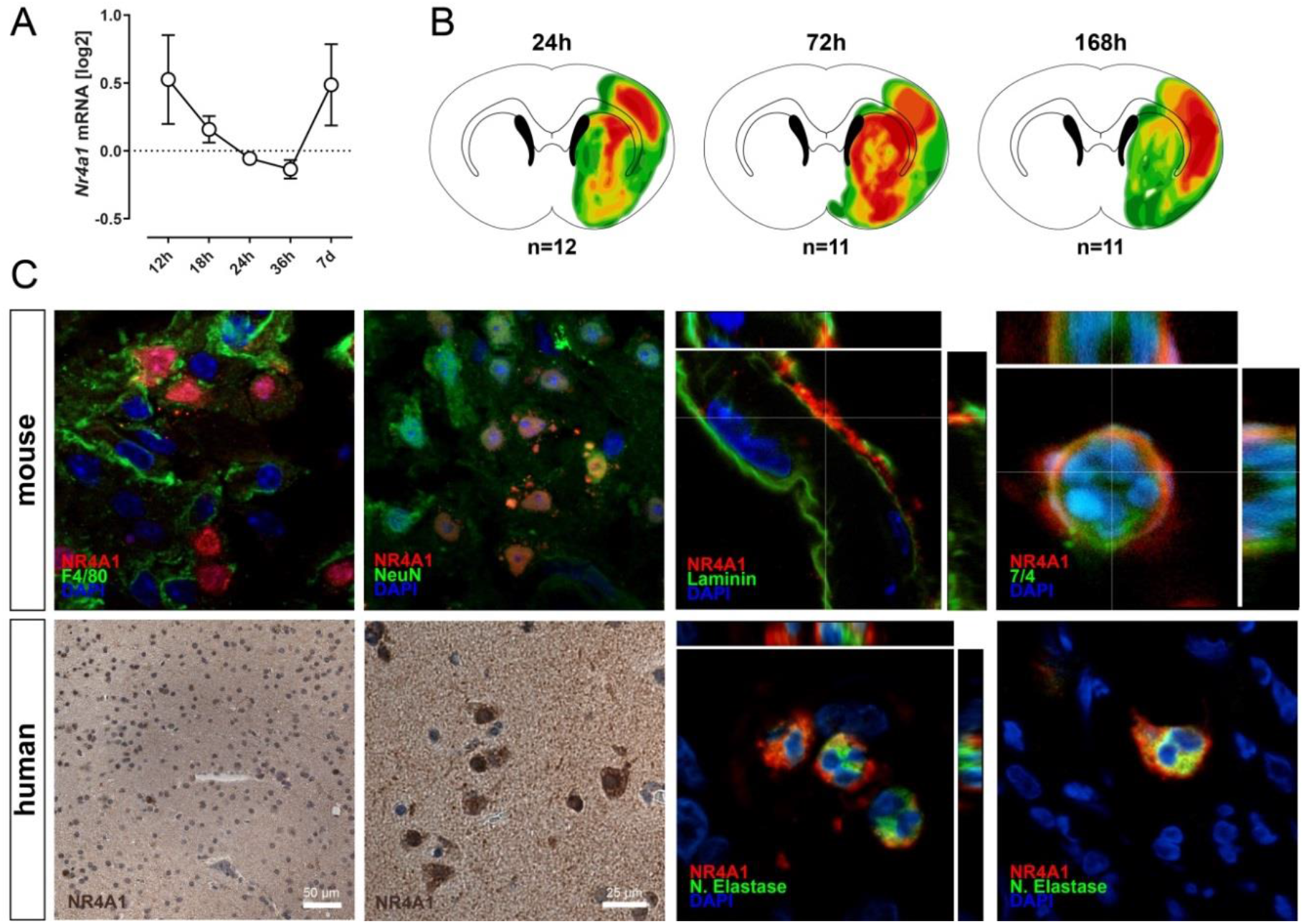
NR4A1 is dynamically expressed following cerebral ischemia. **A.** NR4A1-mRNA expression was evaluated in whole-brain lysates over the time-course of 7 days post MCAO (sham, post 12h, 18h, 24h, 36h and 7d each n=4). **B.** NR4A1-protein signal heatmaps were generated using immunofluorescence stained coronal sections of wildtype mice post 24h (n=12), post 72h (n=11) and post 168h (n=11) after experimental occlusion of the MCA. **C.** Co-localization studies of NR4A1-signal were performed on mice and human stroke specimen. Mouse NR4A1-double staining was done with marker for microglia and monocytes/macrophages (F4/80), neurons (NeuN), blood-vessels (laminin) and neutrophil granulocytes (Ly6B.2 clone 7/4). Staining pattern of 3,3’-Diamino-benzidin labelled NR4A1 indicates particularly neurons and potentially immune cells as NR4A1 source. Immunofluorescence double staining of NR4A1 with neutrophil marker neutrophil elastase confirms neutrophil granulocytes as further NR4A1-expressing cell type.

### NR4A1 limits cerebral ischemia induced brain damage

We next tested a potential role for NR4A1 within stroke pathology using a loss-of-function paradigm. For this purpose, we induced experimental stroke in wildtype (WT) and NR4A1-deficient mice (KO) and compared functional outcome, infarct size, blood-brain barrier damage and induction of pro-inflammatory genes between the two groups. General neuroscore assessment showed a significant recovery decline in the group of NR4A1-deficient mice when compared to the control group (Fig. 2A). Locomotor functional outcome was determined by the rotarod and the foot fault test. Rotarod (Fig. 2B) and foot fault testing (Fig. 2C) confirmed the delayed functional deterioration of NR4A1-deficient mice during the 72h post MCAO investigation period when compared to the wildtype animals. We then extended our investigation to the brain tissue damage progression during the acute phase after MCAO. Twenty-four hours after experimental stroke, no differences in infarct size could be detected between wildtype and NR4A1-deficient mice, matching neuroscore and functional testing results. However, 72h after MCAO, NR4A1-deficient mice showed significantly increased infarct volumes when compared to the wildtype animals (Fig. 2D). Analysis of blood-brain barrier disruption further showed increased IgG-fluorescence signal intensity in NR4A1-deficient mice, confirming an increased brain tissue damage in mice lacking NR4A1 (Fig. 2E). In a similar line, quantitative RT-PCR of whole brain lysates revealed that expression of several proinflammatory genes, including *Ptgs2* (coding COX-2), *Il6* (IL-6) and *Nos2* (iNOS) was significantly increased in whole brain lysates of NR4A1-deficient mice when compared to the wildtype group 72h after MCAO (Fig. 2F), whereas *Tnf* (TNF-α) and *Nfkbia* (IκBα) were not altered. These data suggest that NR4A1 modulates the inflammatory response during the acute phase and thereby restrains the pathophysiological brain damage following stroke.

**Figure 2.**
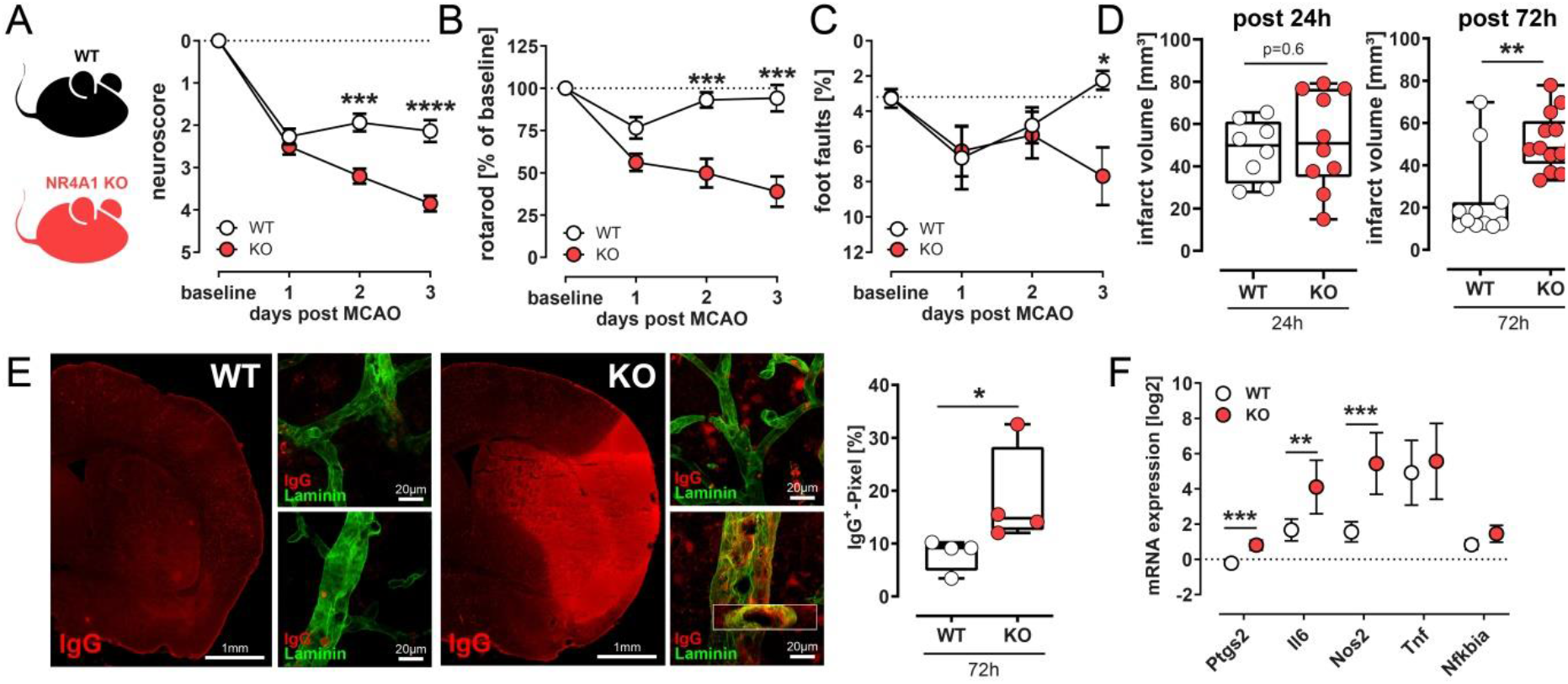
NR4A1 mediates protective effects in cerebral ischemia pathophysiology. **A.** Wildtype and NR4A1-deficient mice were subjected to 30 minutes of MCA occlusion and the neurological deficit score rating was done over the time-course of 3 days post MCAO (WT n=15, KO n=20; ***p<0.001, ****p<0.0001, ANOVA). **B.** Locomotor function was evaluated with the rotarod-test prior to MCAO and on day 1, 2 and 3 post reperfusion (WT n=15, KO n=20; ***p<0.001, ANOVA). **C.** Functional outcome was further assessed using the foot-fault test prior to MCAO and on day 1, 2 and 3 post reperfusion (WT n=14, KO n=17; *p<0.05, ANOVA). **D.** Mean infarct volumes were calculated from 15-20 cryosections (collected at 300 μm intervals) of wildtype and NR4A1-deficient mice at 24h (WT n=8, KO n=10; p=0.6) and at 72h after reperfusion (WT n=11, KO n=13; **p=0.002, *t* test). **E.** Transient MCAO occlusion led to IgG immunoglobulin G extravasation in wildtype and NR4A1-deficient mice. Staining shows co-localization of IgG (red) with endothelial marker pan-Laminin (green). Quantification of IgG-signal intensity suggests an increased damage of the blood-brain barrier in NR4A1-deficient mice (WT n=4, KO n=4; *p=0.029, *t* test). **F.** Expression of pro-inflammatory markers *Ptgs2* (coding COX2), *Il6* (IL-6), *Nos2* (iNOS), *Tnf* (TNF-α) and *Nfkbia* (IκBα) in wildtype (n=5) and NR4A1-deficient (n=5) mice compared to sham operated animals (n=3; **p<0.01, ***p<0.001, *t* test).

### Damage-limiting NR4A1-effects are mediated by neutrophil granulocytes

In order to support our hypothesis that immune cells are responsible for the protective NR4A1 effect in stroke we generated bone-marrow chimeric mice using irradiated wildtype mice (CD45.1) transplanted and reconstituted with either wildtype (congenic CD45.2; WT►WT) or NR4A1-deficient (congenic CD45.2; KO►WT) bone marrow (Fig. 3A). Notably, mice that were transplanted with KO bone marrow cells showed significantly delayed functional deterioration and increased infarct sizes after 30 min of MCAO when compared to the WT that received WT cells (Fig. 3B and C), further indicating a non-brain resident origin of the damage limiting NR4A1-effects. In light of the key role of NR4A1 in T cells in the context of CNS autoimmunity (Liebmann et al., 2018), we hypothesized that the damage limiting NR4A1-effects might be introduced via T cells. However, adoptive transfer of wildtype or NR4A1-deficient T cells into Rag-1-deficient mice did not result in any differences with regard to functional outcome (Fig. 3E) and infarct size development (Fig. 3F), indicating that T cells do not mediate the protective NR4A1-effects in stroke pathology. As innate immune subpopulations become early activated during cerebral ischemia (Beuker et al. 2021), we decided to quantify brain-resident and infiltrating myeloid cells, microglia and neutrophil granulocytes in NR4A1-deficient and wildtype mice after 30 min of MCAO using flow cytometry. We did not detect any differences in the proportions of infiltrating monocytes, macrophages and brain resident myeloid cells (Fig. 3G), whereas the number of infiltrating neutrophil granulocytes was substantially increased in NR4A1-deficient animals when compared to the wildtype mice (Fig. 3H). This could be confirmed by immunofluorescence (Fig. 3I). Although we observed a significant upregulation of neutrophil chemoattractants *Ccl2* and *Cxcl1* (Strecker et al., 2017) in whole brain lysates of wildtype and NR4A1-deficient mice 6h and 24h after onset of ischemia, there was no difference in expression levels between both groups, suggesting that increased neutrophil influx is not related to increased levels of chemoattractants in the brain of NR4A1-deficient mice (Fig. 3J). In order to determine whether neutrophil granulocytes mediate the effects of NR4A1 we depleted Ly6G^+^ neutrophils 24h before stroke induction using a neutrophil-depletion-antibody (Fig. 3K). Importantly, neutrophil depletion abrogated the detrimental effects seen in NR4A1-deficient mice as Ly6G-mAb-treated mice showed full functional recovery (Fig. 3L) and infarct volume (Fig. 3M) comparable to the isotype-treated wildtype group. In conclusion, our data suggest a neutrophil mediated anti-inflammatory and neuroprotective role for the transcription factor NR4A1 after cerebral ischemia.

**Figure 3.**
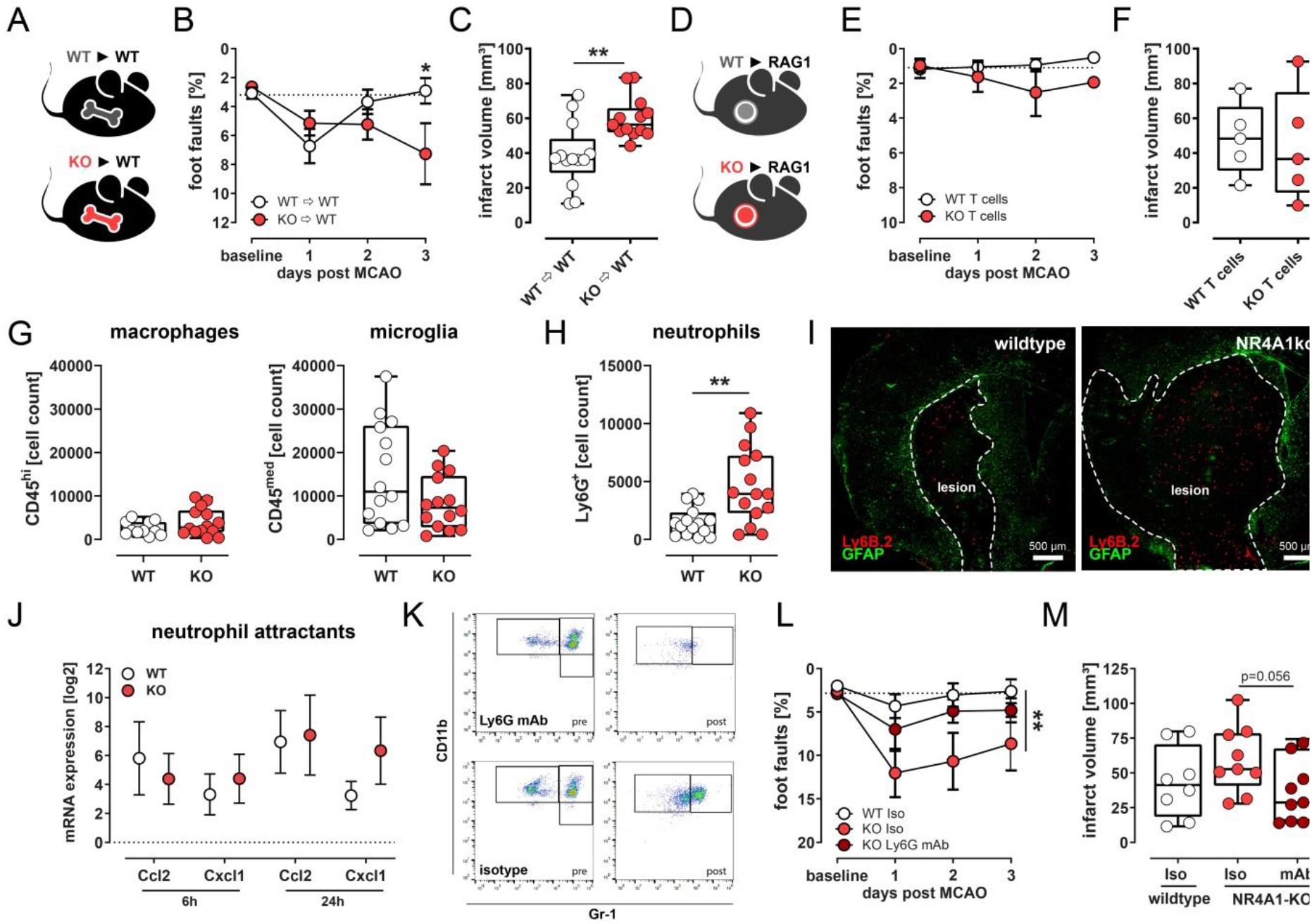
NR4A1 protective effects are mediated by neutrophil granulocytes. **A.** Schematic depiction of the chimerism approach. CD45.1 mice (WT) were sublethally irradiated and 24h after irradiation, animals were either reconstituted with congenic bone marrow wildtype cells (WT►WT; CD45.2) or bone marrow cells of NR4A1-deficient mice (KO►WT; CD45.2). **B.** Foot-Fault tests were performed prior to (baseline) and 24h, 48h, and 72h after MCAO (WT►WT n=13, KO►WT n=13; *p≤0.05, ANOVA). **C.** Mean infarct volumes were calculated from toluidine stained coronal sections in intervals of 300 μm (WT►WT n=13, K►kWT n=13; **p≤0.01, *t* test). **D.** Schematic depiction of adoptive T cell transfer. Either CD3^+^/WT T cells or CD3^+^/NR4A1-deficient T cells were intravenously administered to Rag-1-deficient mice. Twenty-four hours later, mice were subjected to MCAO. **E.** Rag-1-deficient mice were subjected to foot-fault testing prior to and 24h, 48h and 72h after MCAO. **F.** Infarct volumes were compared between Rag-1-deficient mice which received either WT- or NR4A1-deficient T cells. **G.** Cell counts of infiltrating myeloid cells (CD11b^+^/CD45^high^) and microglia (CD11b^+^/CD45^med^), were analyzed in brain tissue isolates by FACS analysis (WT n=14, KO n=14). **H.** FACS analysis of neutrophil cells in brain tissue isolates (WT n=15, KO n=15; **p<0.002, *t* test). **J.** Expression of neutrophil chemokines *Ccl2* and *Cxcl1* was analyzed 6h and 24h after MCAO (WT n=4 (6h) n=5 (24h), KO n=5 (6h) n=4 (24h)). **K.** Representative FACS-plot of blood drawn from mice pre- and post-neutrophil depletion. Mice were treated with either Ly6G antibody (500 μg) or with isotype control IgG (500 μg) 24h prior to MCAO. Shown are CD11b+/Gr-1+-neutrophil counts before and after depletion. **L.** Foot-Faults were recorded prior to and 24h, 48h, 72h post MCAO (WT isotype n=7, KO isotype n=10, KO Ly6G mAb n=12; **p<0.01, ANOVA). **M.** Mean infarct volumes were calculated from coronal brain sections collected at 300 μm intervals (WT isotype n=8, KO isotype n=9, KO Ly6G mAb n=11; KO isotype vs. KO Ly6G mAb, p=0.056, *t* test). For gating strategy of flow cytometry analysis see supplementary figure 2.

### NR4A1 regulates metabolic cell activity in neutrophil granulocytes

We further determined the role of NR4A1 on neutrophil metabolism using wildtype and NR4A1-deficient HoxB8-immortalized neutrophil granulocytes in an *in-vitro* setup. Interestingly, following PMA-stimulation, neutrophils lacking NR4A1 showed an increased basal but attenuated maximal respiration when compared to the wildtype cells suggesting that NR4A1-deficient neutrophils tend to exhaust faster when compared to the wildtype group (Suppl. Fig.3A). In parallel, NR4A1-deficient neutrophils show decreased glycolysis and glycolytic capacity upon stimulation, indicating possible deficiencies in neutrophils, as glycolysis is their major metabolic pathway and critical for important functions such as phagocytosis and neutrophil extracellular trap formation (Suppl. Fig.3B).

### Ligand-mediated activation of NR4A1 improves functional outcome of experimental stroke

In light of the strong modulating role of NR4A1 on stroke outcome, we next asked whether pharmacological targeting of this receptor might represent a therapeutic strategy to limit stroke induced brain damage related to immune cell influx. To this end, wildtype mice were treated with the NR4A1-specific pharmacological ligand Cytosporone-B (Zhang et al., 2008) 3 or 12 hours after reperfusion (Fig. 4A). Notably, Cytosporone-B applied 3 hours after reperfusion resulted in improved functional outcome in the foot fault (Fig. 4B) and the rotarod test (Fig. 4C), and application at 3 and 12 hours after reperfusion further resulted in reduced infarct size when compared to the respective placebo-treated group (Fig. 4D; Suppl. Fig. 1). This protective effect of Cytosporone-B was not observed in NR4A1-deficient mice, thus demonstrating the receptor specificity of the effect (Fig. 4E-H). Taken together, therapeutic targeting of NR4A1 exerts protection in experimental stroke.

**Figure 4.**
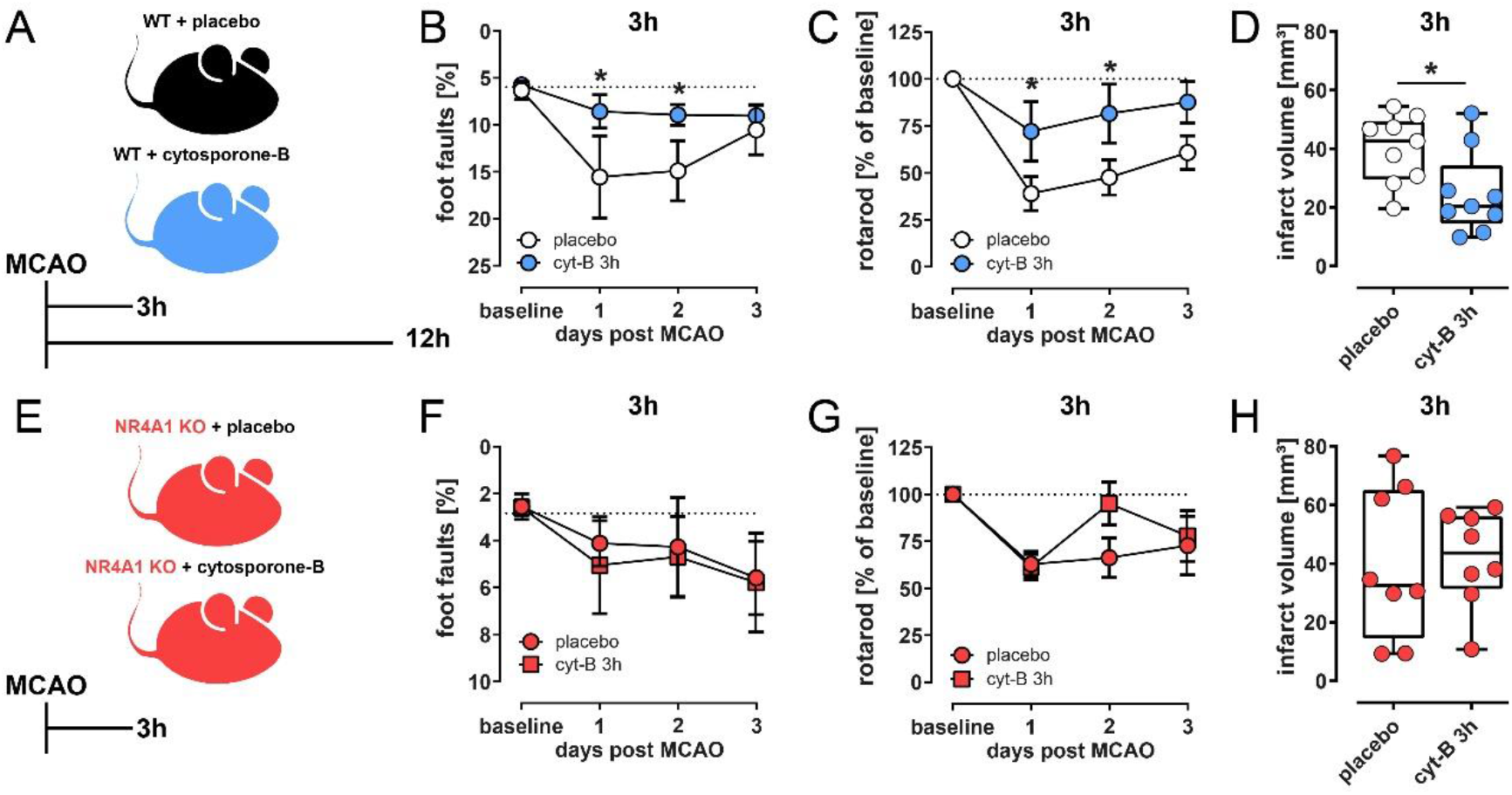
Ligand-mediated activation of NR4A1. **A.** Wildtype mice were treated either with placebo or Cytosporone-B (13 mg/kg body weight) 3h, 6h or 12h after reperfusion. **B.** Functional outcome was evaluated using the foot fault test and the **C.** Rotarod test (placebo n=8-10, Cytosporone-B n=9-10; *p<0.05, AVONA). **D.** Mean infarct volumes calculated from toluidine stained coronal brain sections of mice treated with placebo (n=9) or Cytosporone-B (n=9) 3h after reperfusion (*p<0.05; *t* test). **E.** As further control, NR4A1-deficient mice were treated either with placebo or Cytosporone-B (13 mg/kg body weight) 3h after reperfusion. **F.** Foot-fault testing was performed in NR4A1-deficient mice treated either with placebo or Cytosporone-B (placebo n=6, Cytosporone-B n=5). **G.** Rotarod testing was performed on placebo or Cytosporone-B treated NR4A1-deficient mice over the time-course of 3 days post MCAO (placebo n=8, Cytosporone-B n=8). **H.** Mean infarct volumes were calculated from coronal brain slices of placebo or Cytosporone-B treated mice 3d post MCAO (placebo n=8, Cytosporone-B n=8).

## Discussion

In this study we show that the nuclear receptor NR4A1 plays a significant role in the pathophysiology of ischemic stroke. NR4A1 is expressed within the ischemic mouse and human brain and the lack of NR4A1 results in considerably increased brain damage and deteriorated functional recovery. Therefore, we conclude that NR4A1 exerts damage-limiting effects after cerebral ischemia. Based on experimental evidence from other CNS diseases (Liebmann et al., 2018, Wang et al., 2018), we first assumed T cells as the key mediators of NR4A1-mediated effects in stroke pathophysiology. Indeed, bone-marrow chimera experiments show that the NR4A1-effect is primarily mediated by immune cells of hematogenous origin. However, this effect is not mediated by T cells. While further searching for the potential NR4A1-effect mediating cell population, particularly immigrating neutrophils caught our attention, and indeed, neutrophil depletion abrogated NR4A1-mediated effects in stroke. We therefore conclude that NR4A1 in neutrophils limits the inflammation-induced damage in the context of stroke. Neutrophils have been shown to be involved in a plethora of immune cell driven diseases including ischemic stroke and are further the first immune cell subtype to enter the ischemic brain (Beuker et al., 2021). Recent studies suggest a dual role for distinct neutrophil subtypes and therefore resolving mechanics of neutrophil activation and regulation could lead to the identification of promising new therapeutic approaches (Chen et al., 2021). Numerous studies have already shown that NR4A1 has modulatory effects on disease-mediating cell populations (Carlin et al., 2013; Ipseiz et al., 2014; Liebmann et al., 2018). Hence, NR4A1 might also control neutrophil granulocyte activation. And indeed, in an experimental setup using immortalized neutrophils, we could demonstrate a role for NR4A1 within neutrophil metabolism. Based on these findings we hypothesized that pharmacological ligand-mediated NR4A1-modulation might ameliorate ischemia-induced immune response. Although NR4A1 is a so-called orphan receptor for which no natural ligands have yet been identified, synthetic ligands are available and have recently been successfully used in attenuating pro-inflammatory mediators (Cho et al. 2007, Zhang et al. 2008). Cytosporone-B mediated NR4A1-acitivation has further been shown to ameliorate osteoarthritis by reducing chondrocyte-mediated inflammation (Xiong et al., 2020), to attenuate inflammatory responses in a Parkinson’s disease model (Yan et al., 2020) or even in an experimental model of influenza virus infection via modulation of alveolar macrophages (Egarnes et al., 2017). Indeed, application of Cytosporone-B results in reduced infarcts and improved functional outcome and thus represents a promising future candidate for stroke treatment. A potential weakness of this work could be that neurons and endothel-associated cells also express NR4A1 following stroke. This was not further addressed due to the strong evidence of a neutrophil-mediated effect and should certainly be addressed in future work. However, applying Cytosporone-B in a gain-of-function paradigm gave us the possibility to compare functional outcome, infarct size and immune cell activation directly with the loss-of-function setup used at the start of our study. Therefore, we still consider the results obtained here as a valuable contribution to the understanding on the role of neutrophils in stroke. For ethical issues and to keep mouse numbers low, we decided to test our hypotheses in young male mice to reduce sex related variables. Therefore, the results presented here should be validated in a further study with e.g. in female and in older mice with comorbidities.

The understanding of the role of infiltrating leukocytes within ischemic pathophysiology is of utmost importance for the development of new therapeutic approaches in stroke medicine. In summary, we here show that the nuclear receptor NR4A1 has a substantial role within stroke pathology. The damage limiting characteristics of NR4A1 are mainly neutrophil driven and imported from circulation into the ischemic tissue. Since ligand-mediated NR4A1-activation with Cytosporone-B has been shown to reduce ischemic damage and improve functional outcome in experimental stroke, NR4A1 may therefore represent a potential therapeutic target for stroke treatment. A fine-tuned neutrophil modulation via nuclear receptors could therefore open up promising new possibilities for the development of new treatment strategies.

## Acknowledgements

We thank Maike Hoppen, Birgit Schmeddes, Annika Engbers and Christine Salin for excellent technical assistance. This work was supported by the Deutsche Forschungsgemeinschaft (DFG), grant KL 2199/5-1 and MI 1547/3-1.

## Author contributions

Conception and design of the study: JKS, ML, LK, JM. Providing materials and reagents: TK, CT, HW. Acquisition and analysis of data: JKS, ML, StH, CB, ASP, LK, JM. Drafting manuscript and figures: JKS, ML, LK, JM. Revision and approval of manuscript: all authors.

## Competing interests

The authors declare no competing interests.

## Supplementary figures

**Supplementary Figure 1.**
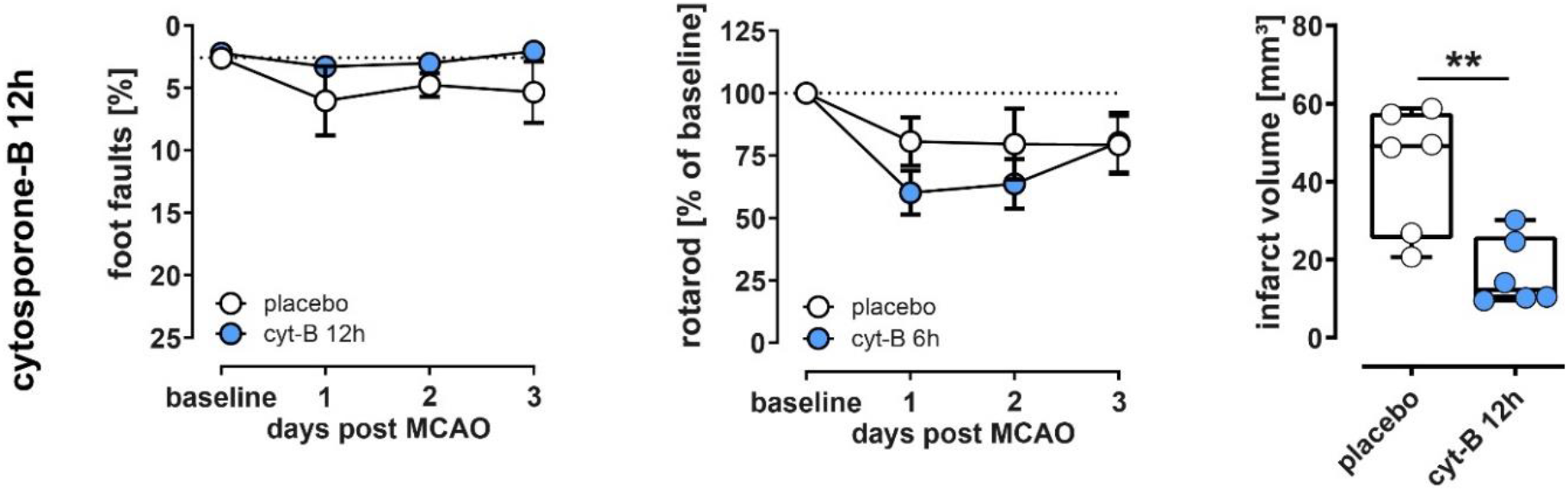
Functional outcome was evaluated using the foot fault test and the rotarod test. Mean infarct volumes were calculated from toluidine-stained coronal brain sections of mice treated with either placebo (n=6-9) or Cytosporone-B (n=6-9) 12h after reperfusion (**p<0.01; *t* test).

**Supplementary Figure 2.**
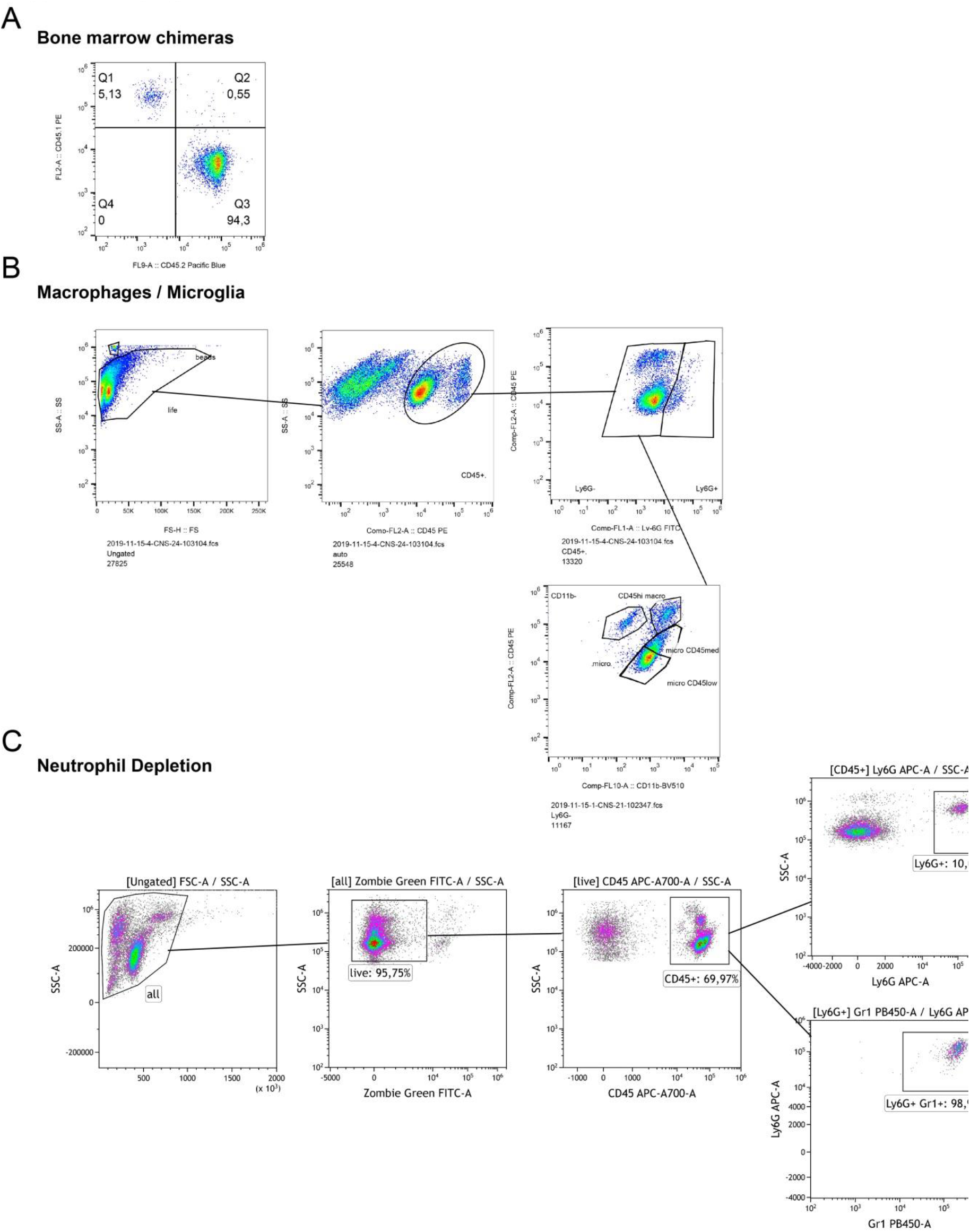
Gating strategy to identify cell types by flow cytometry. **A.** After 8 weeks of reconstitution, chimerism grade was validated by flow cytometric differentiation between the allelic CD45 variants in leukocytes of tail vein blood samples. **B.** Representative gating strategy of the main cell types identified by flow cytometry as depicted in Fig. 3G. **C.** Representative gating strategy of the main cell types identified by flow cytometry as depicted in Fig. K, L and M.

**Supplementary Figure 3.**
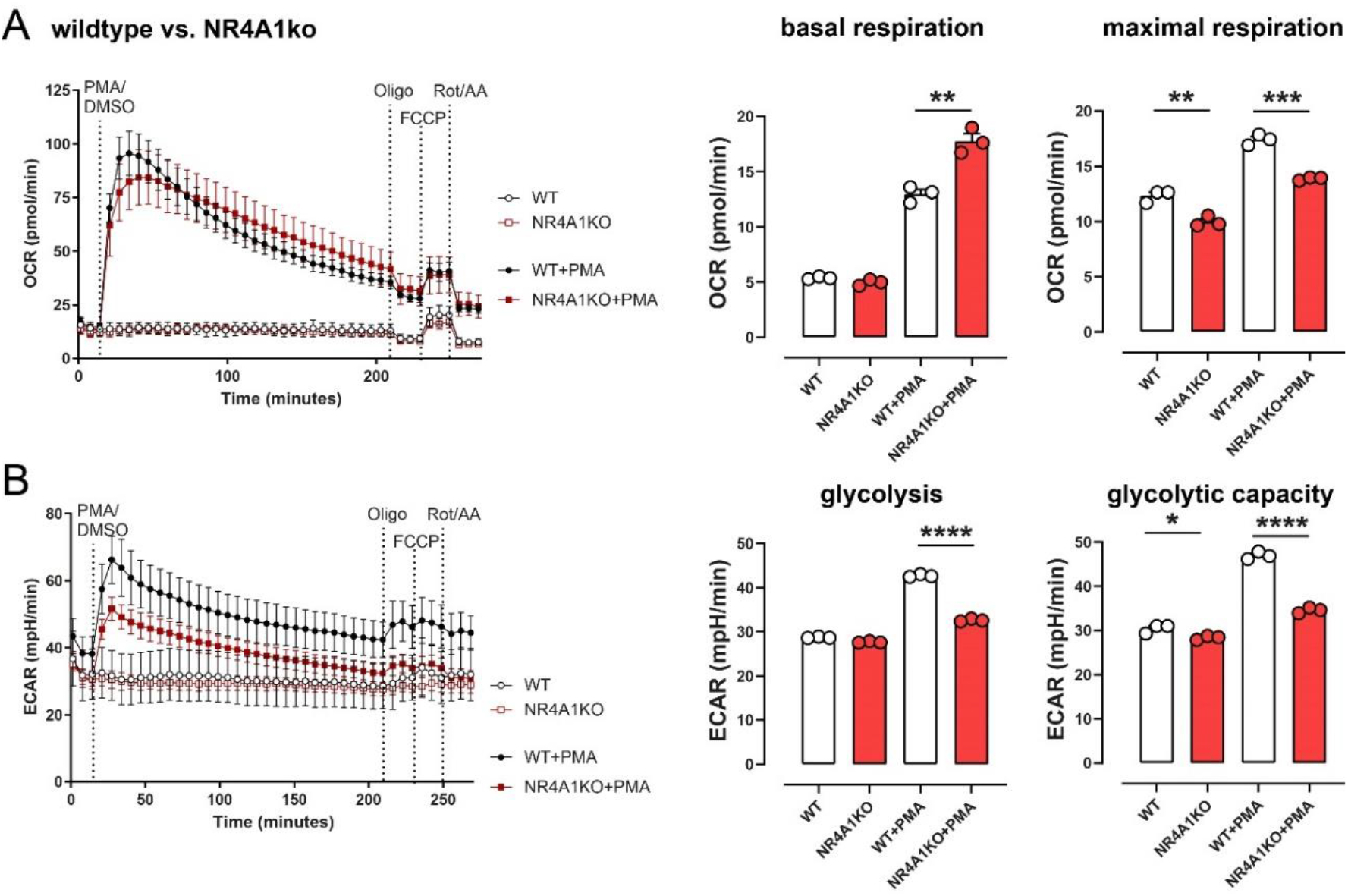
Oxygen consumption rate (OCR) and extracellular acidification rate (ECAR) of wildtype and NR4A1-deficient HoxB8-immortalized neutrophil granulocytes were determined following 5 days of differentiation. Cells were stimulated either with or without 600 nM phorbol-12-myristat-13-acetat (PMA, Sigma-Aldrich) for 3 h. **A.** Graphs depict OCR of the four investigated groups over the time course of 4 hours (left) and mean basal and maximal respiration (right, **p<0.01, ***p<0.001; *t* test). **B.** Graphs depict ECAR of the four investigated groups over the time course of 4 hours (left) and mean glycolysis and glycolytic capacity (right, *p<0.05, ****p<0.0001; *t* test).

**Table S1.**
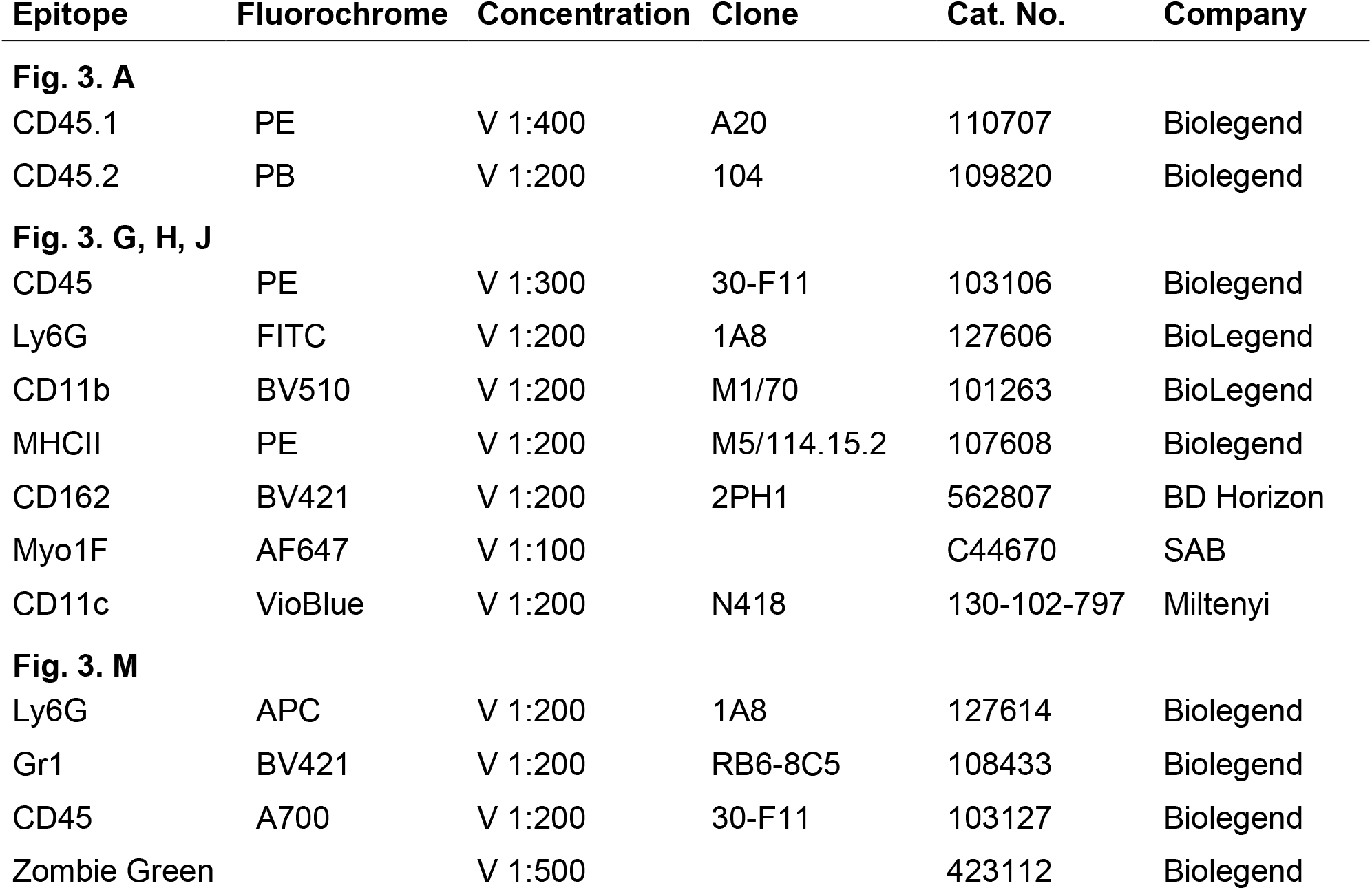
List of murine antibodies used for flow cytometric analysis

## Notes

### Competing Interest Statement

The authors have declared no competing interest.

